# Instructor-learner neural synchronization during elaborated feedback predicts learning transfer

**DOI:** 10.1101/2021.02.28.433286

**Authors:** Yi Zhu, Victoria Leong, Yingying Hou, Dingning Zhang, Yafeng Pan, Yi Hu

## Abstract

The provision of feedback with complex information beyond the correct answer, i.e., elaborated feedback, can powerfully shape learning outcomes such as transfer, i.e., the ability to extend what has been learned in one context to new contexts. However, an understanding of neurocognitive processes of elaborated feedback during instructor-learner interactions remains elusive. Here, a two-person interactive design is used during simultaneous recording of functional near-infrared spectroscopy (fNIRS) signals from adult instructor-learner dyads. Instructors either provided elaborated feedback (i.e., correct answer and an example) or simple feedback (i.e., correct answer only) to learners during a concept learning task. Our results showed that elaborated feedback produced comparable levels of retention to simple feedback, however, transfer was significantly enhanced by elaboration. We also noted significant instructor-learner neural synchronization in frontoparietal regions during the provision of elaborated feedback, especially when examples were provided. Further, interpersonal neural synchronization in the parietal cortex successfully predicted transfer of knowledge to novel contexts. This prediction was retained for both learner-delayed and learner-preceding neural synchronization. These findings point toward transfer effects of elaborated feedback provided in a social context can be predictable through interpersonal neural synchronization, which may hold important implications for real-world learning and pedagogical efficacy.

**Educational Impact and Implications Statement:** Feedback provides learners with crucial information regarding the gap between what has currently been achieved and what remains to be achieved, and thus plays a critical role in any learning process. In real-world settings, feedback is typically provided and received through social interaction, and high-quality “elaborated feedback” contains complex information that goes beyond the correct answer. This study aims to elucidate the neurocognitive processes underpinning elaborated feedback during instructor-learner interactions. We detected significant instructor-learner neural synchronization in mutual frontoparietal brain regions during elaborated feedback, particularly during the provision of specific elaborated information (i.e., concrete examples). Moreover, this synchronization (including learner-delayed and learner-preceded synchronization) in the parietal region predicted whether the learners transferred learning to novel examples of learned psychology concepts. This study advances current understanding on the neural mechanisms for elaborated feedback and the role of social interaction in feedback effects. These results may have important implications for successful real-world learning and communication, and related pedagogical applications in educational settings.

## Introduction

### Learning through social interaction

As we navigate the world, knowledge and skills are often learned on the basis of communication with others during social interaction. The recent decade has witnessed a paradigm shift toward the concurrent measurement of multiple individuals engaging in social interaction (Dai et al., 2018; Kingsbury & Hong, 2020; Redcay and Schilbach, 2019; Schilbach et al., 2013; Wheatley et al., 2019), including infant-adult dyads (Leong et al, 2017; Piazza et al., 2020; Santamaria et al, 2020; Wass et al, 2020) and individuals with neuropsychiatric disorders (Bilek et al, 2017; Leong & Schilbach, 2019). Relevant research has indicated that interpersonal neural synchronization (INS) might underlie social interaction and underpin successful communication (for reviews, see Hasson et al., 2012; Redcay & Schilbach, 2019). For example, Stephens et al. (2010) demonstrated that when communication was successful, the information provider’s brain activity was spatiotemporally coupled with the information receiver’s; INS also showed provider- or receiver-preceding patterns, indicating the provider’s dominance and the receiver’s prediction, respectively.

### Elaborated feedback as a powerful driver in learning

In communication and learning, feedback is a powerful driver of behavioural change as it provides the information regarding the gap between what is achieved and what is aimed to be achieved (Hattie & Timperly, 2007; Mory, 2004). Prior research has identified feedback as a significant factor in student achievement, and learning motivation (e.g., Lepper & Chabay, 1985; Narciss & Huth, 2004). Although it is of great significance, feedback has been regarded as one of the least understood features in the instructional design (Cohen, 1985; Gagne, 1970). In real-world settings, feedback is oftentimes provided and received during two-person interactions, and contains complex information beyond correct answer such as illustrative examples (Hattie & Timperly, 2007). Any type of feedback supplying more complex information than correct answer is generally considered as elaborated feedback (Kulhavy & Stock, 1989). Elaborated feedback has been found to deepen the understanding and promote the transfer to novel contexts (Bangert-Drowns et al., 1991; Butler et al., 2013; Finn et al., 2018; Kulhavy & Stock, 1989, Bransford et al., 1999). However, a scientific understanding of the how elaborated feedback takes effects on learning during social interaction, remains largely elusive.

### Single brain correlates of feedback

Using single-subject experimental designs, a number of studies have established that frontoparietal brain regions including the anterior cingulate cortex (ACC), the dorsolateral prefrontal cortex (DLPFC), and parietal lobules were implicated in the process of feedback messages such as yes-no verification and correct answer, which is regarded as simple feedback (Cavanagh et al., 2011; Crone et al., 2008; Mars et al., 2005; van Duijvenvoorde et al., 2008; Zanolie et al., 2008). Specifically, the ACC was responsible for basic functions such as error detection and expectation violation (Cavanagh et al., 2011; Luft et al., 2013; Mars et al., 2005), while the DLPFC and the superior parietal lobule was engaged in more complex processes such as error correction and performance adjustment (Crone et al., 2008; van Duijvenvoorde et al., 2008; Zanolie et al., 2008). Brain activation in these regions was related to feedback-based learning outcomes such as the memorization of paired-associates (Arbel et al., 2013), response inhibition (McCormick and Telzer, 2018) and performance on reading and mathematics (Peters et al., 2017). To understand more about neurocognitive processes of elaborated feedback during social interaction, the simultaneous investigation of brain signals from interactive dyads is essential but lacking.

### The role of INS in elaborated feedback effects

Within the general domain of social interaction and communication, INS has been found to hold specific implications of effective learning and instruction (Bevilacqua et al., 2018; Dikker et al., 2017; Holper et al., 2013; Meshulam et al., 2021; Nguyen et al., 2020; Pan et al., 2018; 2020; Piazza et al., 2021; Zheng et al., 2018). Based on the simultaneous recording of functional near-infrared spectroscopy (fNIRS) signals from multiple individuals during learning and instruction without the strict restraint of movement (Boas et al., 2014; Pinti et al., 2018), research has identified INS associated with learning outcomes. For instance, INS in the frontal cortex during educational interactions served as a correlate of learners’ performance on singing (Pan et al., 2018) and on statistics (Liu et al., 2019). Besides, instructor-preceding neural synchronization in temporoparietal areas predicted the learners’ performance on numerical reasoning (Zheng et al., 2018). Once feedback is combined with more complex information beyond the correctness, it becomes intertwined with instruction (Hattie & Timperley, 2007). Thence, synchronized brain activity in instructor-learner dyads may offer a new lens into how elaborated feedback takes effects on learning in naturalistic educational settings.

### The present study

Here, we applied fNIRS to simultaneously record brain signals from adult instructors and learners during an ecologically valid yet experimentally controlled educational interaction. Learners studied psychology concepts and received elaborated feedback or simple feedback from instructors. Elaborated feedback contained the correct answer and an example, illustrating the concepts in concrete and real-world situations, while simple feedback only contained the correct answer. Post-learning, learners were assessed for whether they recognized the definitions of learned psychology concepts (i.e., retention measure) and whether they transferred learning to identify novel examples of learned psychology concepts (i.e., transfer measure). We hypothesized that elaborated feedback enhanced learning performance, especially on the transfer measure, relative to simple feedback. Providing and receiving elaborated feedback would synchronize instructor-learner dyads’ brain activity, potentially in frontoparietal regions. Adults rely on the parietal cortex to process the informative and efficient feedback for performance adjustment or error correction (Crone et al., 2008; van Duijvenvoorde et al., 2008). Elaborated feedback, regarded as informative and efficient for the concept learning, facilitates the transfer of knowledge to novel contexts (Butler et al., 2013; Finn et al., 2018). Accordingly, we further hypothesized that parietal instructor-learner neural synchronization would predict learning performance, especially transfer effects.

## Methods

### Ethics statement

This study was carried out according to the guidelines in the Declaration of Helsinki. The study procedure was approved by Human Research Protection Committee at our University. All participants gave their written informed consent prior to the experiment. Participants were financially compensated for their participation.

### Participants

Twenty-four healthy, female, right-handed participants were recruited as instructors. They were required to major in psychology and complete at least one of teacher education courses. Besides, forty-eight healthy, female, right-handed participants were recruited as learners. They were required to not major in psychology. Twelve instructors were randomly assigned into the elaborated feedback group (age *M* = 21.75, *SD* = 2.42), while the other twelve into the simple feedback group (age *M* = 21.25, *SD* = 2.93, *t*_(22)_ = 0.46, *p* = 0.65). Each instructor was randomly paired with up to two learners. The instructor taught each of the two learners using the same type of feedback (either elaborated or simple feedback) individually over two adjacent days, resulting in a between-subject design for both leaners and instructors. We chose this design to blind instructors (all psychology majors) to the experimental purpose and achieve higher consistency in task delivery across learners. Accordingly, 48 dyads composed of one instructor and one learner were formed. The age of learners did not differ between the elaborated feedback group (*M* = 19.63, *SD* = 1.95) and simple feedback group (*M* = 19.79, *SD* = 1.77, *t*_(46)_ = 0.31, *p* = 0.76). We merely recruited female dyads to control for the potential impacts of gender difference (Baker et al., 2016; Cheng et al., 2015; see also Hu et al., 2018; Pan et al., 2018; 2020 for similar settings). All participants were naïve with respect to the purpose of the study.

### Materials

Materials used for instruction and learning were about a set of ten psychology concepts from the topic of judgement and decision making (Rawson et al., 2015). Each concept has a term, a one-sentence definition and two examples (view details in Table S1). Examples illustrated target concepts in concrete and real-world situations. Examples used in the current study were adapted from psychology textbooks (Hou, 2018; Pastorino & Doyle-Portillo, 2008; Zimbardo et al., 2012) and materials used by previous studies on feedback-based learning (Finn et al., 2018; Rawson et al., 2015). The specific use of materials was described together with the experimental procedures as follows.

### Experimental protocol

The experiment was carried out over two visits to the laboratory, with the interval of one or two days (Figure 1a).

**Figure 1.**
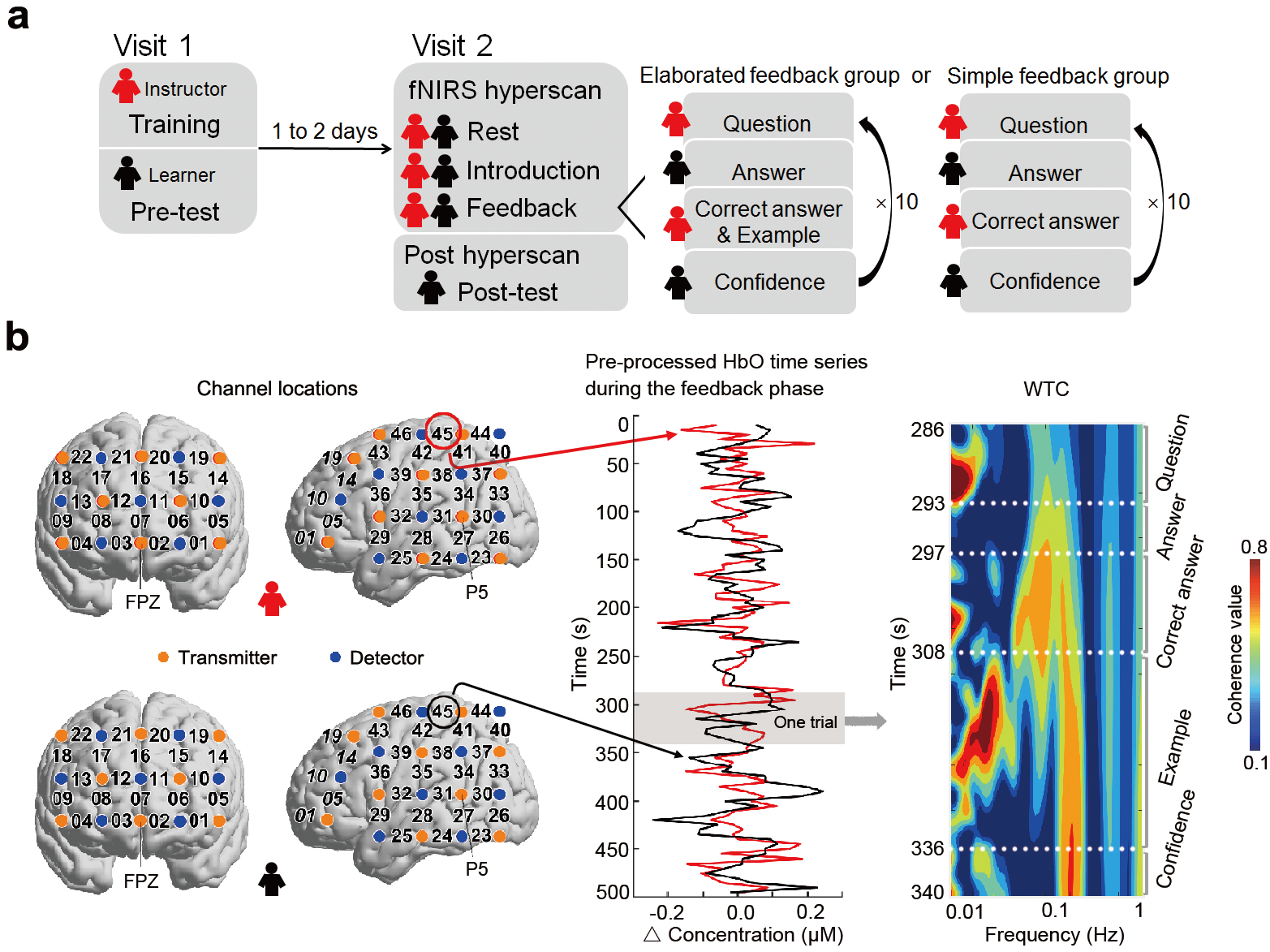
Experimental protocol, channel locations and WTC analysis. *Note.* (**a**) Schematic of the experimental protocol. During the first visit, instructors underwent a standardized training on the instructional procedure and content and leaners completed a pre-learning test. During the second visit, instructor-learner dyads first rested. Then instructors introduced 10 concepts, during which the term and definition were orally presented twice. Next, learners re-studied 10 concepts one by one based on instructors’ feedback (simple feedback of correct answer only or elaborated feedback of correct answer and example). Their brain activity was simultaneously recorded via fNIRS. Post hyperscanning, learners completed a post-learning test assessing both knowledge retention and knowledge transfer. (**b**) Locations of measurement channels and illustration of WTC analysis. On the left panel, two optode probes were placed over instructors’ and learners’ frontal and left temporoparietal areas, respectively. Measurement channels were located between one transmitter (orange) and one adjacent detector (blue). Location references were placed at FPZ and P5 according to 10-10 international system. On the middle panel, sample data were one instructor-learner dyad’s preprocessed HbO time series from CH45 during the feedback phase. On the right panel, the resulting WTC matrix (frequency × time) corresponding to one trial was visualized with color bar denoting the values. HbO, oxy-hemoglobin; WTC, wavelet transform coherence.

During visit 1, learners completed a pre-learning test (< 15 min) assessing their prior knowledge relative to those ten psychology concepts. Specifically, learners were required to match 10 definitions with 10 terms from provided 12 terms (c.f. Allen and Brooks, 1991; Finn et al., 2018; Murphy, 2004). The extra two terms were also from the same topic of judgement and decision making (view details in Table S1). The prior knowledge was quantified in forms of accuracy on pre-learning test (i.e., dividing the number of correctly matched concepts by the number of all concepts). As expected, learners had comparable prior knowledge in the elaborated vs. simple feedback group (*M* ± *SD*, 0.58 ± 0.19 vs. 0.58 ± 0.26, *t*_(46)_ = 0, *p* > 0.999). Besides, learners completed a battery of scales with regard to learning and motivation: (*i*) Achievement Goal Orientation (Button et al., 1996); (*ii*) Academic Self-efficacy (Pintrich & Groot, 1990); (*iii*) Learning Engagement (Schaufeli et al., 2002). No significant differences on scales for two feedback groups were detected (*t*s < 1.60, *p*s > 0.10). During visit 1, instructors underwent a standardized training on the instructional procedure and content (~ 30 min). Afterwards, instructors brought home the print copies of the instruction materials and were required to learn and recite the concepts for their definitions and examples (see details in Table S1) at home. Upon coming back to the laboratory for visit 2, instructors were required to correctly recall the instructional procedure, together with the definitions and examples of two randomly selected concepts by the experimenter. Instructors were not allowed to carry out formal instruction until they met those requirements.

Visit 2 consisted of two sessions: fNIRS hyperscanning and post-hyperscanning. During the first session, instructors and learners sat face-to-face approximately 1 meter apart, wearing the fNIRS equipment. This session consisted of three phases: rest, introduction and feedback.

In the rest phase (300 s), both instructors and learners kept their eyes closed, motion restrained and mind relaxed. In the introduction phase, instructors introduced 10 concepts one by one with the term and definition orally presented twice. The introduction order of the concepts was self-decided by instructors in advance. In this phase, learners listened to the introduction with the permission of requesting the repetition of unclear parts. This phase was self-paced and instructor-learner dyads in elaborated vs. simple feedback group spent comparable time (337.77 s ± 62.02 vs. 330.78 s ± 66.86, *t*_(46)_ = 0.38, *p* = 0.71).

In the feedback phase, learners re-studied the 10 concepts based on the instructor’s feedback. The flow relevant to one concept, i.e., one trial, could be split into four periods: question, answer, feedback and confidence. Specifically, instructors first presented a definition and questioned learners which term corresponded to the definition. Then, learners gave an answer. Next, instructors provided elaborated or simple feedback to learners depending on which feedback group she was assigned in. Simple feedback merely involved the correct answer, which consisted of the term and the definition, while elaborated feedback involved the correct answer and an additional example. Finally, learners judged the confidence that they would correctly answer the relevant questions in the post-hyperscanning session via number keyboards (0–9, *very low* to *very high*). One trial for elaborated feedback group was exemplified as follows.

Instructor: The tendency, once an event has occurred, to overestimate one’s ability to have foreseen the outcome. Which term did this definition correspond to?
Learner: Hindsight bias.
Instructor: The correct term is hindsight bias, whose definition is the tendency, once an event has occurred, to overestimate one’s ability to have foreseen the outcome. Here is an example. Some students will pat the thighs after the teacher announces the correct answer and say “I know this is the choice!”
Learner: (press one number).

In this phase, the order of 10 concepts was also self-decided by instructors in advance, but should be different from that in the introduction phase. As expected, instructor-learner dyads in elaborated vs. simple feedback group spent longer time in the feedback period (339.54 s ± 48.42 vs. 137.13 s ± 28.38, *t*_(46)_ = 17.67, *p* < 0.001). To note, instructor-learner dyads in elaborated feedback group spent 136.04 s ± 22.22 and 203.50 s ± 30.06 for the correct answer and example part, respectively. The whole process of the fNIRS hyperscanning session was also recorded via a digital video camera (Sony, HDR-XR100, Sony Corporation, Tokyo, Japan).

Following the feedback phase, the fNIRS hyperscanning device was immediately unequipped and participants completed a scale assessing task load (Hart, 2006), which showed no difference between the two feedback groups (*t* = 0.82, *p* = 0.421). Next, learners completed a post-learning test (< 15 min) measured both the retention of knowledge and the transfer of knowledge to novel contexts. On the retention measure, learners were required to match 10 definitions with 10 terms from provided 12 terms, which was identical with the pre-learning test. On the transfer measure, learners had to match 10 novel examples with 10 terms from provided 12 terms (c.f. Finn et al., 2018). To note, the selection of examples for use in elaborated feedback (i.e., Example 1 in Table S1) vs. transfer measure (i.e., Example 2 in Table S1) was previously decided by the experimenters without replacement. The elaboration example and the specific context/topic provided for the transfer measure were not similar as assessed by an additional group of raters (*N* = 20, 16 females, age *M* = 24.45, *SD* = 2.89; see Supplementary Methods for details).

### fNIRS data acquisition and preprocessing

Instructors’ and learners’ brain activity was simultaneously recorded during the hyperscanning session of visit 2 using an ETG-7100 optical topography system (Hitachi Medical Corporation, Japan). Two optode probes were used for each participant: a 3×5 probe covering frontal areas (eight transmitters and seven detectors resulting in 22 measurement channels, i.e., CH1–22) and a 4×4 probe covering left temporoparietal areas (eight transmitters and eight detectors resulting in 24 measurement channels, i.e., CH23–46), see Figure 1b for the reference and channel locations. The probes were placed over frontal and temporoparietal areas because these regions have been implicated in feedback-based learning (Crone et al., 2008; Luft, 2014; van Duijvenvoorde et al., 2008) as well as learning and instruction (Liu et al., 2019; Pan, et al., 2018; Zheng et al., 2018). Temporoparietal areas were focused on the left hemisphere rather than the right hemisphere due to the former is dominant for language functions (Ojemann et al., 1989; Vigneau et al., 2006), which is an essential component of concept learning. The correspondence between NIRS channels and measured points on the cerebral cortex was determined using the virtual registration approach (Singh et al., 2005; Tsuzuki et al., 2007; see details in Table S2).

The optical data were collected at the wavelengths of 695 and 830 nm, with a sampling rate of 10 Hz. The preprocessing of fNIRS data was performed using custom MATLAB (MathWorks, Natick, MA, USA) scripts and Homer2 toolbox (version 2.2, Huppert et al., 2009). The raw optical intensity data series were first converted into changes in optical density (OD). Channels with very low or high OD, which exceeded 5 SDs, were marked as unusable and removed from the analysis. Next, OD time series were screened and corrected for motion artifacts using a channel-by-channel wavelet-based method. The Daubechies 5 (db5) wavelet was chosen (Molavi & Dumont, 2012) and the tuning parameter was set to 0.1 (Cooper et al., 2012). A band-pass filter with cut-off frequencies of 0.01–1 Hz was applied to the OD data in order to reduce the slow drift and high frequency noise. The OD time data were then converted into oxyhemoglobin (HbO) and Deoxyhemoglobin (HbR) concentration changes based on the modifier Beer-Lambert Law (Cope & Delpy, 1988). In the current study, we mainly focused on HbO concentration change, which was considered as an indicator of the change in regional cerebral blood flow with higher signal-to-noise ratio (Hoshi, 2007) and has been more widely used in fNIRS hyperscanning research (e.g., Cheng et al., 2015; Hu et al., 2017; Jiang et al., 2015; Pan et al., 2017; Dai et al., 2018; Yang et al., 2020).

### Data analysis

#### Behavioral data analysis

Learning performance was assessed by post-learning test and quantified in forms of accuracy (i.e., dividing the number of correctly answered items by the number of all items). Besides, learners’ knowledge immediately before feedback (i.e., on the answer period of the feedback phase) was also quantified in forms of accuracy, which was comparable between simple feedback group (*M* ± *SD*, 0.67 ± 0.21) and elaborated feedback group (0.62 ± 0.15, *t*_(46)_ = 0.82, *p* = 0.41).

First, we sought to verify whether conceptual knowledge was promoted by elaborated feedback. Because each instructor was randomly assigned to teach two learners, learners were nested within instructors. A linear mixed model (West et al., 2014) was thus fitted on learners’ accuracy including fixed effects of test time (pre-learning vs. post-learning), plus random effects on learner and instructor identity. Accuracy on the answer period of the feedback phase and the duration of elaborated feedback were additionally entered in the model to control for their potential effects.

Next, we investigated whether elaborated feedback promoted the learning relative to simple feedback. A linear mixed model was fitted on learners’ accuracy on the retention measure, including a fixed effect of feedback type (elaborated vs. simple), plus random effects of learner and instructor identity. Accuracy on the pre-learning test, accuracy on the answer period of feedback phase and the duration of feedback were additionally entered in the model to control for their potential effects. Besides, a parallel model was fitted on learners’ accuracy on the transfer measure.

Finally, an additional linear mixed model was conducted on confidence ratings including a fixed effect of feedback type (elaborated vs. simple), plus random effects of learner and instructor identity.

All behavioral analyses were computed using functions implemented in MATLAB (R2018a, MathWorks). Linear mixed models were constructed using *fitlme* function. Restricted maximum likelihood was used to estimate the models. *F* and *p* values were derived using *anova* function based on Satterthwaite approximation.

#### fNIRS data analyses

##### WTC analysis

Interpersonal neural synchronization (INS) between instructors and learners was computed by a wavelet transform coherence (WTC) algorithm, which estimates the correlation of a pair of time series as a function of frequency and time (Grinsted et al., 2004; Torrence & Compo, 1998). First, preprocessed HbO time series were extracted from homologous regions (following previous studies, e.g., Cui et al., 2012; Hu et al., 2018; Jiang et al., 2012; Liu et al., 2019; Pan et al., 2018; 2020). For instance, two signals (*i* and *j*) could be respectively extracted from instructors’ CH45 and the learners’ CH45 (Figure 1b). Then, WTC of signals was computed by following formula:

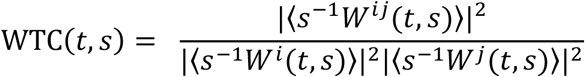

where *t* denotes the time, *s* indicates the wavelet scale, 〈·〉 represents a smoothing operation in time and scale, and *W* is the continuous wavelet transform. Then, a 2-D (time × frequency) WTC matrix was generated (Figure 1b, see more details in Chang & Glover, 2010; Grinsted et al., 2004).

In this study, we specifically investigated INS associated with elaborated feedback (for general instruction and learning, see Liu et al., 2019; Pan et al., 2018; 2020; Zhang et al., 2018). To this end, time points corresponding to the start and the end of feedback (i.e., the feedback period, Figure 1b) were marked based on the recorded videos and was adjusted for the delay-to-peak effect by 6 s (Cui et al., 2009; Jiang et al., 2015). Accordingly, elaborated feedback could be further segmented into two parts (i.e., correct answer and example, Figure 1b).

##### Cluster-based permutation test

Interpersonal interactions as opposed to resting state elicited significantly larger INS (Cui et al., 2012; Jiang et al., 2012). For each dyad and each channel combination, WTC values during the feedback period and the rest phase (leaving out first and last minutes to retain more steady data) were respectively time-averaged, and then converted into Fisher *z*-values. Accordingly, we sought to identify frequency-channel clusters showing significantly larger WTC during elaborated feedback vs. rest using a cluster-based permutation test. It is a non-parametric statistical test that offers a solution to the problem of multiple comparisons for multi-channel and multi-frequency data (Maris & Oostenveld, 2007). We conducted it following five steps. First, we ran frequency-by-frequency and channel-by-channel linear mixed models including a fixed effect of task (feedback vs. rest), plus random effects of learner and instructor identity. Considering the process of elaborated feedback was self-paced, duration was entered in the model to control for its potential effect. Next was to identify channels (46 in total) and frequency bins (80 in total, ranging from 0.01 to 1 Hz), at which the task effect was significant (feedback > rest, *p* < 0.05). To note, we excluded 12 respiration-related frequency bins from 0.15 to 0.3 Hz and 7 cardiac-related frequency bins above 0.7 Hz (Nozawa et al., 2016; Zheng et al., 2018), remaining 60 frequency bins (see in Supplementary material, Figure S1). Third was to form clusters composed of neighboring channels (≥ 2) and neighboring frequency bins (≥ 2) and compute the statistic for each cluster by summing all *F* values. Fourth, repeat WTC analysis and the first step using permuted data and calculate the statistics for each cluster identified in the third step for 1000 times. The permutation was conducted by randomly pairing one learner’s dataset with another instructor’s dataset. As the length of datasets varied across dyads, the longer dataset was trimmed to the same length as the shorter one for each random pair (Reindl et al., 2018), see details in the Supplementary Materials and Figure S2. Finally, the observed cluster statistics were compared with the results of 1000 permutations (both converted to square roots to normalize the distributions) with *p* value assessed by following formula (Theiler et al., 1992):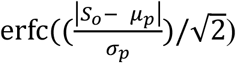, *S*_o_ denotesobservedclusterstatistic, *μ*_*p*_, *σ*_*p*_ respectively denote the mean and standard deviation of permutation results. The clusters with *p* value < 0.05 were regarded as significant. Besides for elaborated feedback, the cluster-based permutation test was also conducted on each of two parts of elaborated feedback, i.e., correct answer and example, and simple feedback, i.e., correct answer only, respectively.

##### Contrast analysis

To further characterize brain regions more strongly synchronized by different forms of feedback information (example vs. correct answer), a contrast analysis was performed on the significant clusters identified by the cluster-based permutation test. To control for individual differences, we used clusters’ ΔWTC in the following analyses, which was computed by subtracting WTC (averaged by channels and frequency bins contained in the cluster) during task from that during rest, and then converted into Fisher *z*-values. Before entering the contrast analysis, time series of ΔWTC during elaborated feedback was segmented into two parts, i.e., correct answer and example, based on the recorded videos (Figure S3). Instructor-learner dyads in the elaborated feedback group spent 136.04 s ± 22.22 and 203.50 s ± 30.06 for the correct answer and example part, respectively (*t* = 15.58, *p* < 0.001). Then the contrast between different forms of feedback information was conducted following two steps (Figure S3). First, compare ΔWTC during correct answer and example contained in elaborated feedback. Specifically, a linear mixed model was fit on ΔWTC associated with two parts of elaborated feedback, including a fixed effect of feedback information (example vs. correct answer), as well as random effects of learner and instructor identity. Considering the varying data length across feedback information and across dyads, duration of feedback information was entered in the model to control for its potential effect. Second, compare ΔWTC during simple feedback (correct answer only) and the example part of elaborated feedback, using an identical linear mixed model as that in the first step. Multiple comparisons were corrected using the false discovery rate (FDR) method (Benjamini and Hochberg, 1995) to calculate *corrected p* values.

#### Behavior-brain relation analyses

Next, we tested whether instructor-learner neural synchronization associated with elaborated feedback predicted learning performance. To control for individual differences, relative accuracy was used in the following analysis, which was computed by subtracting z-score of accuracy on the pre-learning test from that on the post-learning test. A machine learning algorithm, i.e., linear support vector regression (SVR), was applied to train ΔWTC for each identified cluster for the prediction of relative accuracy. To avoid the potential information loss by the trial-averaged ΔWTC value, we instead extracted trial-by-trial ΔWTC values, which was then used as up to ten features for the training. We used a leave-one-out cross-validation approach via Regression Learner APP implemented in MATLAB (R2018a, MathWorks). The prediction analysis was performed by doing such a training first on all but one dyad and then testing on the left-out dyad to examining the generalization of prediction of relative accuracy based on trial-by-trial ΔWTC. The prediction analysis was performed *n* times (*n* = total number of dyads). Prediction accuracy was quantified by the Pearson correlation coefficient (*r*) between the observed and predicted relative accuracy (Hou et al., 2020; Kosinski et al., 2013). The value of *r* ranges from −1 to 1, indicating the worst to best prediction accuracy, with the value of *p* indicating the significance. Considering elaborated feedback unfolded over time, when the aforementioned prediction analyses showed significant results (*r* > 0 and *p* < 0.05), we added various time shifts (instructor’s brain activity was shifted forward or backward relative to the learner’s by 1–14 s, step = 1 s) to the re-computation of prediction analyses, with FDR method (Benjamini and Hochberg, 1995) calculating *corrected p* values.

## Results

### Elaborated feedback promoted the transfer of knowledge

As expected, accuracy on the post-learning test (*M* ± *SD*, 0.83 ± 0.13) was significantly higher than that on the pre-learning test (0.58 ± 0.19, *F*_(1, 23)_ = 58.50, *p* < 0.001, *β* = 0.25, SE = 0.03, 95% confidence interval (CI) = 0.19 to 0.32). It was indicated that elaborated feedback promoted learners’ conceptual knowledge. Next, we investigated whether elaborated feedback relative to simple feedback promoted learning. On the retention measure, learners’ accuracy was comparable in the elaborated feedback group (0.96 ± 0.09) and simple feedback group (0.94 ± 0.14, *F*_(1, 21.17)_ = 1.90, *p* = 0.183, *β* = 0.04, SE = 0.03, 95% CI = −0.02 to 0.09). However, on the transfer measure, a parallel model analysis revealed that learners’ accuracy in the elaborated feedback group (0.70 ± 0.21) was significantly higher than that in the simple feedback group (0.59 ± 0.21, *F*_(1, 15.63)_ = 5.42, *p* = 0.031, *β* = 0.14, SE = 0.06, 95% CI = 0.02 to 0.26). It was indicated that elaborated feedback relative to simple feedback promoted transfer rather than retention of knowledge. Besides, for the confidence rating, no significant effect was revealed (*F*_(1, 22)_ = 0.49, *p* > 0.100).

### Elaborated feedback synchronized instructor-learner dyads’ neural activity in the frontoparietal regions

We investigated whether instructor-learner dyads providing and receiving elaborated feedback as opposed to resting elicited significantly larger WTC using a cluster-based permutation test. Two significant channel-frequency clusters were identified (Figure 2 and Table S3). Cluster 1 was composed of 2 spatially neighboring channels, i.e., CH42, CH45, in 8 frequency bins, ranging from 0.017 to 0.025 Hz (cluster statistic = 11.54, *p* < 0.001). The channels contained in Cluster 1 were approximately located at the left parietal cortex, including the postcentral gyrus (PoCG) and superior parietal gyrus (SPG). Cluster 2 was composed of 3 spatially neighboring channels, i.e., CH05, CH06, CH10, in 7 frequency bins, ranging from 0.017 to 0.024 Hz (cluster statistic = 6.62, *p* = 0.005). The channels contained in Cluster 2 were approximately located at the left frontal cortex, including the superior frontal gyrus (SFG) and middle frontal gyrus (MFG). In addition, instructor-learner synchronization on Cluster 1 and Cluster 2 exhibited temporal patterns, i.e., the learners’ brain activity synchronized with instructors’ with some delay or the opposite (see details in Supplementary Results, Figure S4).

**Figure 2.**
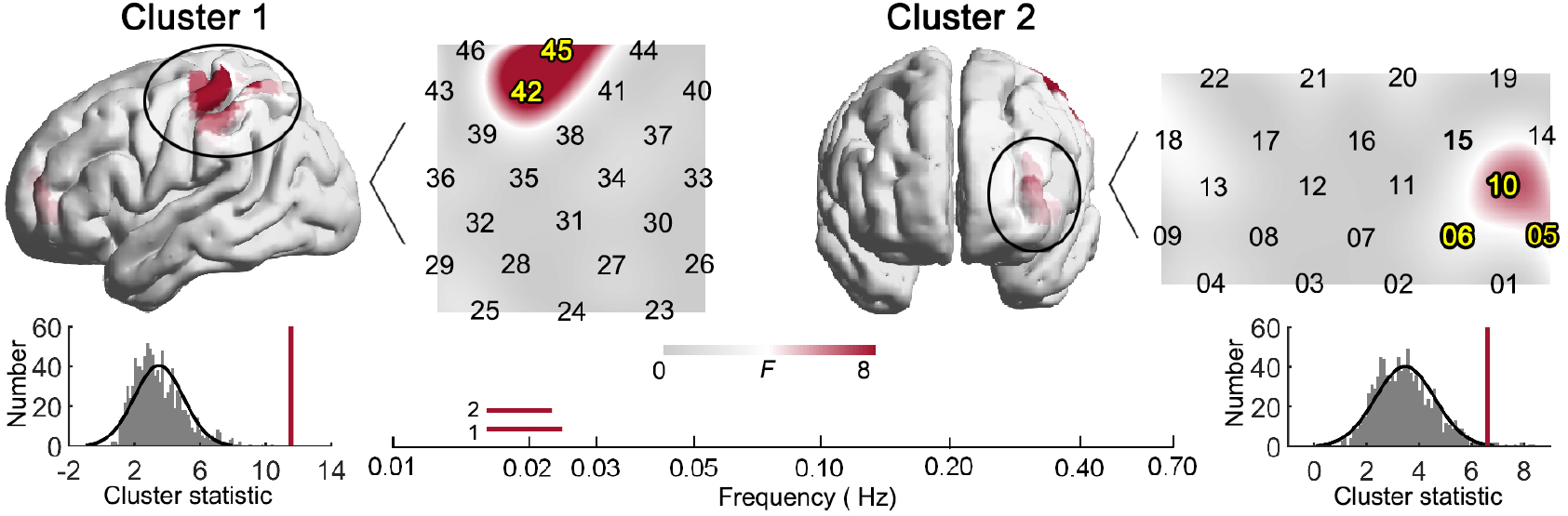
Instructor-learner neural synchronization during elaborated feedback. *Note.* Two significant clusters were identified. Cluster 1 was approximately located at the left PoCG and left SPG within 0.017–0.025 Hz and Cluster 2 was approximately located at the left SFG and left MFG within 0.017–0.024 Hz (with permutation tests, *p*s < 0.001). Spatial locations of the clusters are visualized at a representative frequency bin of 0.02 Hz. Yellow numbers denote channels contained in the clusters. Red horizontal lines denote the frequency bands. Gray histograms depict the frequent distribution of null cluster statistics, while red vertical lines denote observed cluster statistics.

Additionally, granger causality analysis was performed to explore the information flow during the period of elaborated feedback from instructor to learner or from learner to instructor on brain regions corresponding to the identified clusters (see more details in Supplementary Methods). Granger causality analysis revealed significant and comparable bidirectional information flow between the instructor and the learner when providing and receiving elaborated feedback (see more details in Supplementary Results, Figure S2).

### Frontoparietal instructor-learner synchronization was specific to examples

To further characterize the brain regions synchronized by different feedback information, brain activity during elaborated feedback was segmented into two parts (i.e., example and correct answer) and respectively compared with that during resting using a cluster-based permutation test. For the example part of elaborated feedback, two significant channel-frequency clusters were identified (Figure 3 and Table S4). Cluster 3 was composed of 2 spatially neighboring channels, i.e., CH42, CH45, in 8 frequency bins, ranging from 0.018 to 0.027 Hz (cluster statistic = 13.69, *p* < 0.001). The channels contained in Cluster 3 were approximately located at the left parietal cortex, including the PoCG and SPG. Cluster 4 was composed of 3 spatially neighboring channels, i.e., CH05, CH06, CH10, in 8 frequency bins, ranging from 0.015 to 0.023 Hz (cluster statistic = 10.61, *p* < 0.001). The channels contained in Cluster 4 were approximately located at the left frontal cortex, including the SFG and MFG. To note, Cluster 1 and Cluster 3 contained identical channels, while Cluster 2 and Cluster 4 contained identical channels. In addition, the synchronized brain activity on Cluster 3 and Cluster 4 exhibited temporal patterns, i.e., the learners’ brain activity synchronized with instructors’ with some delay or the opposite (see details in Supplementary Results, Figure S4). However, for the correct answer part of elaborated feedback, no significant channel-frequency cluster was identified (Table S4). Simple feedback (only containing the information of correct answer) was also compared with rest using a cluster-based permutation test and no significant channel-frequency cluster was identified (Table S5). It was indicated that instructor-learner neural synchronization on frontoparietal regions was specific to the example rather than correct answer part of elaborated feedback.

**Figure 3.**
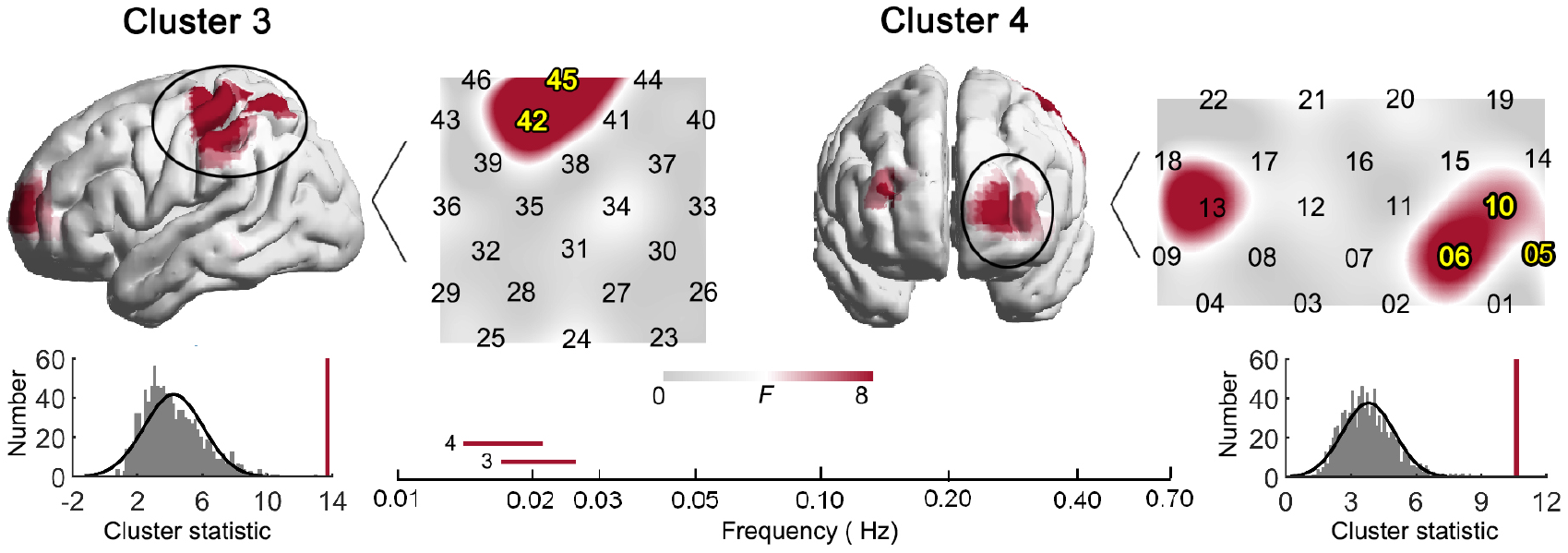
Instructor-learner neural synchronization during the example part of elaborated feedback. *Note.* Two significant clusters were identified. Cluster 3 was approximately located at the left PoCG and left SPG within 0.018–0.027 Hz and Cluster 2 was approximately located at the left SFG and MFG within 0.015–0.023 Hz (with permutation tests, *p*s < 0.001). Spatial locations of the clusters are visualized at a representative frequency bin of 0.02 Hz. Yellow numbers denote channels contained in clusters. Red horizontal lines denote frequency bands. Gray histograms depict the frequent distribution of null cluster statistics, while red vertical lines denote observed cluster statistics.

Next, contrast analysis was conducted between different forms of feedback information (example vs. correct answer) by two steps, on Cluster 3 and Cluster 4, respectively. The first was to compare ΔWTC during the example and correct answer contained in elaborated feedback, and the second was to compare ΔWTC during the example part of elaborated feedback and simple feedback (correct answer only) based on linear mixed models. On Cluster 3, providing and receiving the example vs. correct answer part of elaborated feedback elicited larger ΔWTC (feedback minus rest) (0.10 ± 0.12 vs. 0.09 ± 0.11, *F*_(1, 23.70)_ = 8.21, *p* = 0.009, *corrected p* = 0.018, *β* = 0.15, SE = 0.05, 95% CI = 0.04 to 0.25, Figure 4a), with the duration of feedback information showing a significant effect (*F*_(1, 27.87)_ = 11.486, *p* = 0.002, *β* = −0.002, SE = 0.001, 95% CI = −0.003 to −0.001); providing and receiving the example part of elaborated feedback vs. simple feedback also elicited larger ΔWTC (0.10 ± 0.12 vs. 0.01 ± 0.14, *F*_(1, 26.60)_ = 4.75, *p* = 0.037, *corrected p* = 0.049, *β* = 0.13, SE = 0.06, 95% CI = 0.01 to 0.24, Figure 4a), with the duration of feedback information showing non-significant effect (*F*_(1, 31.17)_ = 0.56, *p* = 0.461, *β* = −0.000, SE = 0.001, 95% CI = −0.002 to 0.001). On Cluster 4, providing and receiving the example vs. the correct answer part of elaborated feedback elicited comparable ΔWTC (0.12 ± 0.13 vs. 0.11 ± 0.13, *F*_(1, 19.73)_ = 2.46, *p* = 0.133, *corrected p* = 0.133, *β* = 0.09, SE = 0.06, 95% CI = −0.03 to 0.22, Figure 4b), with the duration of feedback information showing non-significant effect (*F*_(1, 23.88)_ = 3.48, *p* = 0.074, *β* = −0.001, SE = 0.001, 95% CI = −0.003 to 0.000); providing and receiving the example part of elaborated feedback vs. simple feedback elicited larger ΔWTC (0.12 ± 0.13 vs. 0.03 ± 0.17, *F*_(1, 45)_ = 9.39, *p* = 0.004, *corrected p* = 0.016, *β* = 0.20, SE = 0.06, 95% CI = 0.07 to 0.32, Figure 4b), with the duration of feedback information showing significant effect (*F*_(1, 45)_ = 4.63, *p* = 0.037, *β* = −0.002, SE = 0.001, 95% CI = −0.003 to −0.000).

**Figure 4.**
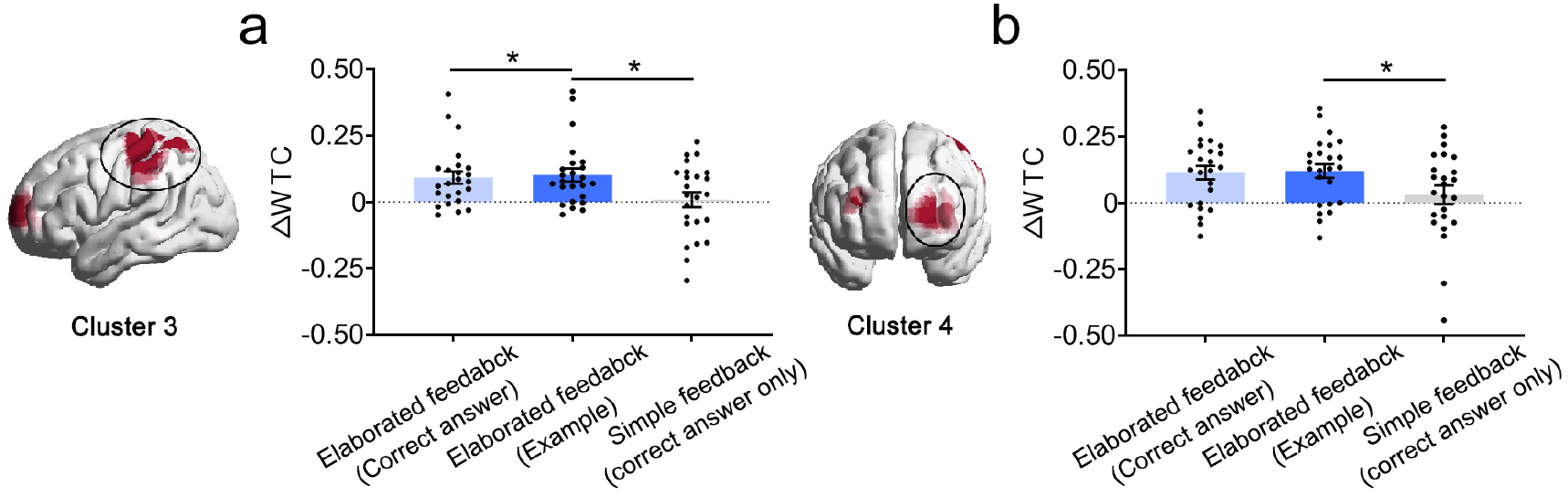
Instructor-learner neural synchronization during example vs. correct answer. *Note.* (**a**) On Cluster 3, example relative to correct answer part of elaborated feedback and simple feedback elicited significantly larger ΔWTC. (**b**) On Cluster 4, example relative to correct answer part of elaborated feedback elicited comparable ΔWTC, while example part of elaborated feedback relative to simple feedback elicited larger ΔWTC. \**p* < 0.05.

### Parietal instructor-learner neural synchronization predicted the transfer of knowledge

Next, we tested whether instructor-learner neural synchronization during providing and receiving elaborated feedback could predict learning performance. A SVR was trained on ΔWTC associated with the example part of elaborated feedback on Cluster 3 and Cluster 4 to respectively predict learners’ accuracy on the post-learning test relative to the pre-learning test. It was revealed in Figure 5a that trial-by-trial ΔWTC on Cluster 3 could successfully predict out-of-sample learners’ relative accuracy on the transfer measure (*r* = 0.57, *R*^2 =^ 32.49%, *p* = 0.004) but not on the retention measure (*r* = 0.25, *R*^2 =^ 6.25%, *p* = 0.241); trial-by-trial ΔWTC on Cluster 4 could not predict learning performance (*r*s < −0.09, *R*^2^s < 0.81%, *p*s > 0.05). A similar prediction pattern was seen for synchronized neural activity associated with elaborated feedback (see more details in Supplementary Results, Figure S6a).

**Figure 5.**
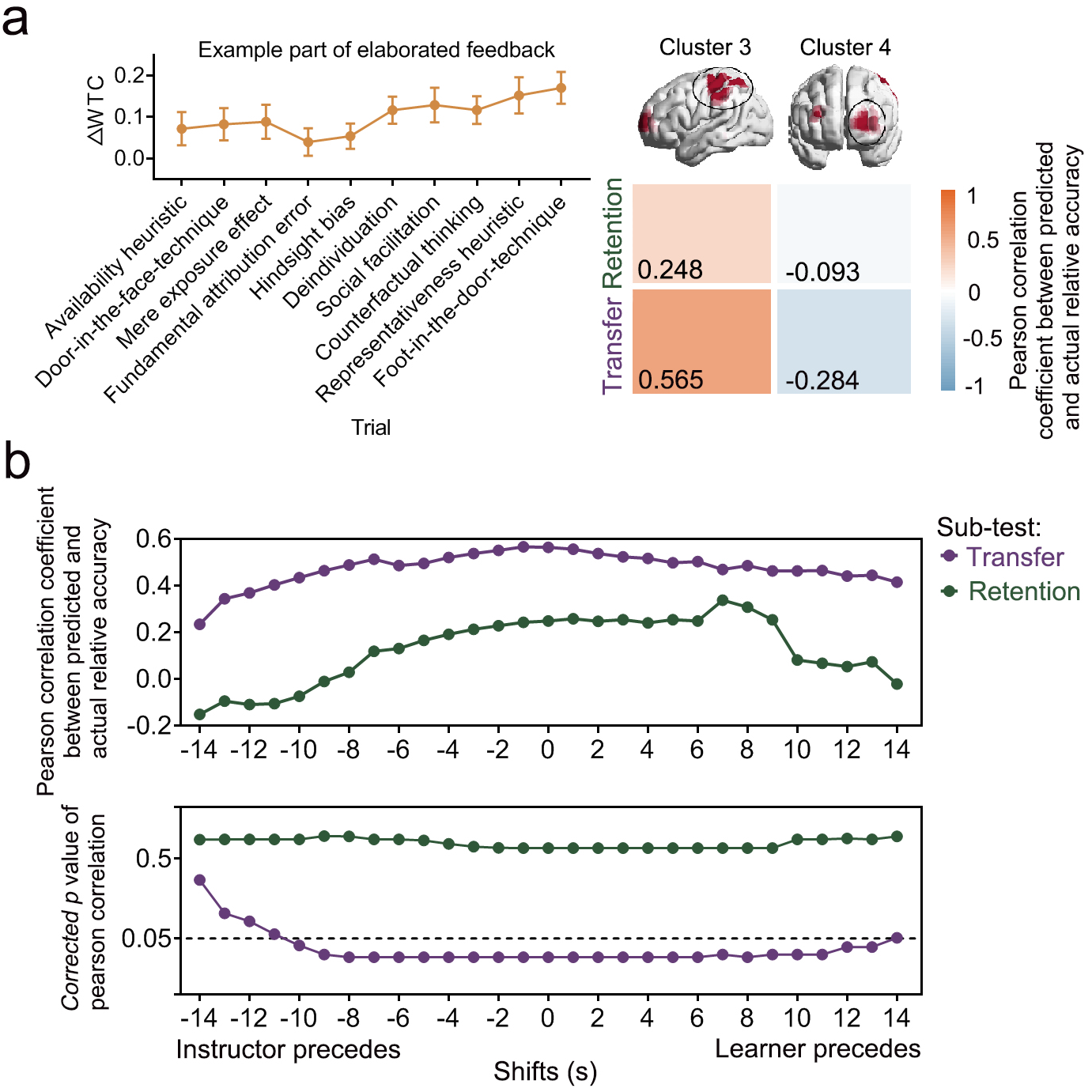
Instructor-learner neural synchronization during the example part of elaborated feedback predicts transfer. *Note.* (**a**) Trial-by-trial ΔWTC on Cluster 3 could successfully predict out-of-sample learners’ relative accuracy on the transfer measure but not on the retention measure. Warmer colors indicate relatively higher prediction accuracy for a given cluster; cooler colors indicate relatively lower prediction accuracy for a given cluster. (**b**). The prediction accuracy for Cluster 3 on the transfer measure was significant when instructors’ brain activity preceded learners’ by 1–10 s and when learners’ brain activity preceded instructors’ by 1–13 s (−10–13, purple).

Moreover, when time shifts were added to re-perform the prediction analysis based on trial-by-trial ΔWTC associated with the example part of elaborated feedback on Cluster 3, the prediction accuracy on the transfer measure was significant when instructors’ brain activity preceded learners’ by 1–10 s and when learners’ preceded the instructors’ by 1–13 s (*corrected p*s < 0.05, Figure 5b). With time shifts, the prediction accuracy on the retention measure remained insignificant (*corrected p*s > 0.05, Figure 5b). With time shifts, a similar prediction pattern was seen for synchronized brain activity associated with elaborated feedback (see more details in Supplementary Results, Figure. S6b).

## Discussion

Our findings support the notion that providing learners with elaborated feedback relative to simple feedback promotes the transfer of conceptual knowledge to novel contexts. The neurocognitive processes of elaborated feedback during instructor-learner interactions were investigated from an inter-brain perspective. When elaborated feedback unfolded overtime, we found synchronized instructor-learner dyads’ brain activity in frontoparietal regions, including the superior frontal gyrus (SFG), middle frontal gyrus (MFG), postcentral gyrus (PoCG) and superior parietal gyrus (SPG). Such instructor-learner synchronization was specific to complex information, i.e., example, contained in the elaborated feedback. Based on a machine learning algorithm, instructor-learner synchronization associated with example in the parietal cortex successfully predicted out-of-sample learners’ ability to transfer knowledge to novel contexts. Such a prediction was retained when instructors’ brain activity preceded learners’ by 1–10 s and when learners’ preceded instructors’ by 1–13s.

Although elaborated feedback is theorized to increase the probability of error correction and the depth of knowledge comprehension (Jacoby et al., 2005; Morris et al., 1977; Tulving & Thompson, 1973), previous studies have demonstrated divergent evidence on its specific effects on learning. For example, compared with correct answer feedback, adding example or explanation to feedback promotes the learning of conceptual knowledge (for both knowledge retention and transfer, Finn et al., 2018; for knowledge transfer only, Butler et al., 2013). However, no greater effects of elaborated feedback relative to correct answer feedback on learning have also been reported (e.g., Andre & Thieman, 1998; Kulhavy et al., 1985; Mandernach, 2005). It may be due to that the added information is too lengthy or complex to be processed and even offsets the effects of correct answer (Kulhavy et al., 1985; Shute, 2008). The present study found that providing learners with elaborated feedback containing example relative to correct answer feedback resulted in comparable retention of knowledge. However, when learners’ ability to transfer conceptual knowledge to novel contexts was tested, elaborated feedback tended to be of benefit. These findings supported the superior effect of elaborated feedback on knowledge transfer rather than knowledge retention. To note, in the current study, learning gains were measured almost immediately after the hyperscanning session. Follow-up studies should have another post-test with a delay interval (e.g., one week) to explore whether the effects of elaborated feedback are retained over longer intervals.

Metacognitive effects of elaborated feedback are also recognized as a crucial factor in feedback research. Correct answer feedback not only facilitates the correction of erroneous responses with high confidence (Butterfield & Metcalfe, 2001, 2006; Pashler et al., 2005), but also calibrates metacognitive errors on low-confidence correct responses (Butler et al., 2008; Thomas & McDaniel, 2013). Feedback, especially elaborated feedback, may improve the calibration and item-level accuracy of metacognitive judgments. In particular, the processing of examples contained in elaborated feedback might affirm or trigger re-evaluation of the learner’s deeper conceptual understanding. Moreover, elaborated feedback provided in a social context involves social cues and its efficacy would be expected to be moderated by social effects such as relationship between the instructor and the learner. Besides, patterns of neural synchronization might differ based on whether participant’s answer in the feedback phase was correct or incorrect. Unfortunately, the limited number of items (only 10) in this study restricted item-level analyses or conditional analyses on correct vs. incorrect responses. Future research is required to explore whether feedback on correct vs. incorrect answers, high vs. low confidence correct answers, or high vs. low confidence errors differs with respect to the sequencing of learner-instructor synchronization (that is, learner-delayed or learner-preceded neural synchronization).

When instructor-learner dyads providing and receiving elaborated feedback, we found synchronized brain activity in frontoparietal regions. Frontoparietal regions such as the anterior cingulate cortex (ACC), DLPFC and parietal lobules are well-localized by single-brain imaging research on feedback-based learning (Cavanagh et al., 2011; Crone et al., 2008; Luft et al., 2013; Mars et al., 2005; van Duijvenvoorde et al., 2008; Zanolie et al., 2008). Activity generated in the ACC, tracks a basic feedback function of error detection and conflict monitoring (Cavanagh et al., 2011; Luft et al., 2013; Mars et al., 2005). Moreover, the DLPFC and parietal lobules play essential role in error correction and performance adjustment (Zanolie et al., 2008; van Duijvenvoorde et al., 2008). Besides, DLPFC is also implicated in social interaction (Kanske et al., 2015; Schurz et al., 2014). In the current study, synchronized brain activity observed approximately in the SFG, MFG, PoCG and SPL, which were spatially proximal to well-defined feedback sensitive regions, may underlie the providing and receiving elaborated feedback by instructor-learner dyads in real-world educational settings. In our study, we further demonstrated that instructor-learner synchronization in frontoparietal regions was specifically associated with complex information, i.e., example, contained in the elaborated feedback, whereas providing and receiving the correct answer failed to synchronize brain activity from instructors and learners. These results suggest that feedback information beyond the correct answer recruit separable brain activity in instructor-learner dyads, which potentially supports the superior effect of elaborated feedback on learning.

Furthermore, based on linear SVR, instructor-learner synchronization associated with example in the parietal cortex rather than frontal regions successfully predicted out-of-sample learners’ ability to transfer knowledge to novel contexts. In comparison with the ACC, parietal lobules mature late in feedback processing (Peters et al., 2016). Adults rely more on the parietal cortex than the ACC to process informative and efficient feedback to adjust performance or correct errors (Crone et al., 2008; van Duijvenvoorde et al., 2008; Zanolie et al., 2008), which plays a more critical role in knowledge acquisition. Concrete examples contained in elaborated feedback tended to be informative and efficient for concept learning and had advantages in facilitating transfer (Bangert-Drowns et al., 1991; Butler et al., 2013; Finn et al., 2018; Kulhavy & Stock, 1989). The current study observed instructor-learner neural synchronization in frontal regions but such neural synchronization had no connection to learning performance. In line with previous research, feedback information tended to activate frontal brain regions (Cavanagh et al., 2011; Mars et al., 2005). However, due to the limited depth of NIR light penetration (Ferrari & Quaresima, 2012), brain activity generated as deep as from the “feedback-related” ACC (Cavanagh et al., 2011) might not have been reliably tracked. Future studies could use fMRI hyperscanning to assess the involvement of INS in frontal regions in feedback-based learning. In this study, whether INS serves as a mechanism that supports learning or it is simply an epiphenomenon also requires further careful and detailed examination (Hamilton, 2021; Wass et al., 2020; Novembre & Iannetti, 2021; Pan et al., 2021a). One way to test the causal role of INS in learning is using a multi-brain stimulation protocol (Novembre et al., 2017; Novembre & Iannetti, 2021; Pan et al., 2021b).

Interestingly, prediction effect of instructor-learner synchronization associated with example in the parietal cortex retained when instructors’ brain activity preceded learners’ by 1–10 s and when learners’ preceded instructors’ by 1–13 s. The processing of high-level linguistic structures such as sentences and paragraphs is at timescale of seconds, whereas that of sound-level acoustic features is milliseconds (Hasson et al., 2015). In average, each example was presented with 2.4 sentences, lasting for about 20.3 second. Therefore, the maximal temporal shifts are more likely to reflect sentence-level rather than word- or syllable-level processing. Transfer tends to occur when the prior learned knowledge is represented at deeper levels, e.g., abstract structure and personal interpretation, instead of surface levels, e.g., specific words and syntax (Graesser et al., 1997; Kintsch, 1998). To extract the abstract structure of knowledge demands a sufficient amount of information being transmitted from instructors to learners and the integration of such information over a time window (Stephens et al., 2010; Tatler et al., 2003). Accordingly, this predicts that learner-delayed neural synchronization may predict transfer effects. If knowledge was represented into personal interpretation, learners would be able to predict the upcoming information before it was completely provided (DeVault et al., 2011; Pickering & Garrod, 2013), resulting in learner-preceding neural synchronization that predicts transfer effects. In the current study, we found that instructor-learner neural synchronization with temporal shifts (both learner-delayed and learner-preceded) could successfully predict transfer, which provides preliminary supporting evidence to the notion that deeper-level representations of knowledge in parietal regions may promote transfer. Nevertheless, as previous research has found that abstract knowledge structure (also called “schema”) is associated with mPFC function (Gilboa, 2017), other brain regions may also play a critical role in deep-level knowledge representations. Future research should specifically address underlying cognitive processes supporting the transfer effect of elaborated feedback by experimental manipulation. To note, the broad significant time window detected in the current study might indicate a lack of temporal sensitivity in blood flow changes to cognitive events (Huppert et al., 2006; Pinti et al., 2020). Considering the broad time window, specific conclusions regarding the directionality of effects may not be drawn.

In current study, several questions deserve noting. First, instructor-learner dyads in the elaborated feedback group spent extra ~200 seconds than those in the simple feedback group during task. The amount of social interaction in dyads might have influenced the synchronization of instructor-learner brain activity (Zheng et al., 2018). Though our linear mixed models controlled for the factor of duration of feedback, it would be ideal for future studies to have a third control group that received simple feedback with time on task equated with the elaborated feedback condition. Second, in accordance with previous hyperscanning studies of educational interactions (Holper et al., 2013; Liu et al., 2019; Pan et al., 2018; 2020), we mainly focused on INS between the instructors’ and learners’ homologous regions across different time lags (i.e., one’s brain activity precedes that of the other). Considering the instructors and the learners are expected to have different roles (i.e., teaching and learning), neural synchronization between different brain regions or that with time lags is expected (Jiang et al., 2021; Zheng et al., 2018; Liu et al., 2017). Due to the limited channels of fNIRS, our optode probe set only covered the frontal cortex and left temporoparietal regions, leaving the functions of other regions unexplored. Future studies are encouraged to consolidate our findings by using whole-brain coverage and by further exploring the neural synchronization between different regions in instructors and learners. Third, frequencies of instructor-learner neural synchronization associated with elaborated feedback were roughly identified within 0.01 to 0.03 Hz, overlapping some of those identified by previous fNIRS hyperscanning studies using communication paradigms (e.g., Jiang et al., 2012; 2015) and education tasks (e.g., Zheng et al., 2018). Future research may wish to further characterize INS for its potential significance in the frequency domain as EEG signals in terms of ranges and functions (Henry, 2006; Teplan, 2002). Fourth, considering that feedback effects could be mediated by learners’ prior knowledge (Fyfe et al., 2012; Krause et al., 2009) and metacognitive judgment (Butler et al., 2008; Thomas & McDaniel, 2013), future work is expected to be more prudent when screening learners. For example, apart from not being Psychology majors, learners are also expected to not have taken a Psychology class in recent years. Their degree of confidence or certainty in the correctness of the testing items should also be assessed. Besides, only female dyads were tested in order to reduce the sample variability, in accordance with previous evidence and recommendations (Baker et al., 2016; Cheng et al., 2015; Tang et al., 2019). Future studies should consolidate and generalize the current findings to male participants. Last but not the least, the critical role of social factors, such as communication mode (e.g., human-human, human-computer) and relationship between instructors and learners (e.g., trust, rapport), in shaping learning from feedback might be a fruitful direction for future investigations.

In summary, the current results suggest that the feedback information beyond the correct answer could promote learning, especially for transfer of knowledge to novel contexts. Extending previous findings based on computer-controlled paradigms, this study used an ecologically valid yet experimentally controlled feedback-based concept learning task carried out by instructor-learner dyads with their brain activity simultaneously measured using fNIRS. As feedback information unfolded over time, instructor-learner neural synchronization was observed in frontoparietal regions, especially when examples were provided, and predicted the transfer of conceptual knowledge to novel contexts. Inter-brain dynamics may provide a novel lens for people to understand more about how elaborated feedback and learner-instructor interactions shape learning and transfer, thence unmasks the neurocognitive basis of feedback provided in a social context and contributes to pedagogical efficacy.

## Notes

**Author Note** Data collection and preliminary analysis were sponsored by the National Natural Science Foundation of China (31872783 and 71942001). We have no conflicts of interest to disclose.

### Competing Interest Statement

The authors have declared no competing interest.

