## Supplementary material for "Instructor-learner neural synchronization during elaborated feedback predicts learning transfer"

### Supplementary Methods

#### The relationship between feedback example and test example

The similarity of the event/context described in the feedback example (i.e., Example 1 in Table S1) and that in the test example (i.e., Example 2 in Table S1) for each of ten concepts was rated (1–7, extremely low to extremely high) by another group of Psychology majors ($N$ = 20, 16 females, age $M$ = 24.45, $SD$ = 2.89). The average rate was 4.21 ± 0.88, varying from 2.85 to 5.25, being not significantly higher than the moderate level (i.e., 4) ($p$ = 0.278), using a Wilcoxon signed-rank test. The rater reliability based on the Kendall coefficient of concordance was 0.313 ($p < 0.001$), indicating the ratings by this group of participants were significantly concordant.

#### Pseudo instructor-learner dyads

The permutation was conducted by randomly pairing the instructor and the learner from different original real dyads under the same condition as a pseudo dyad (Figure S2a). To note, the whole length of data collected in the hyperscanning session was composed of three phases, i.e., rest, introduction and feedback. The rest phase was fixed to last 5 min, while the introduction phase and the feedback phase were both self-paced with varying duration across dyads. The time spent in the feedback phase by 24 real dyads in elaborated feedback group varied from 373.50 s to 898.50 s, with average of 600.34 s and standard deviation of 117.43. To handle the varying length of datasets across real dyads, the longer dataset was trimmed to the same length as the shorter one in each pseudo dyad (Reindl et al., 2018) following two steps (Figure S2b). The first step was to align the starting point of feedback phase by cropping the longer introduction data (i.e., from Learner 3), for which is not focused to be analyzed in the current study. The second step was to align the ending point of feedback phase by cropping the longer feedback data (i.e., from Instructor 1). The approach of pair-randomization would possibly form 552 (24*23) different pseudo instructor-learner dyads. When trimming longer data to match the shorter data from pseudo instructor-learner dyads, the length of cropped data during the feedback phase varied from 1.40 s to 281.80 s, with average of 83.12 s and standard deviation of 62.30.

If necessary, the data separation of the instructors specifically giving simple feedback, elaborated feedback, correct answer and example part of elaborated feedback was based on the marks (based on the recorded videos) from the data with shorter-length feedback phase (i.e., from Instructor 1).

**Instructor-learner neural synchronization with time shifts**

Considering feedback unfolded over time and the participants in this study are expected to have different roles (i.e., instructors and learners), neural synchronization with time shifts (i.e., the learners' brain activity synchronized with instructors' with some delay or the opposite) is expected (Jiang et al., 2012; Liu et al., 2017; Stephens et al., 2010; Zheng et al., 2018). Accordingly, for clusters identified by the aforementioned cluster-based permutation tests, time shifts were added to compare the WTC during elaborated feedback or example part of elaborated feedback vs. resting. Specifically, the HbO time series of the instructor's brain activity was first shifted forward or backward relative to the learner's by 1–14 s (step = 1 s). Next, for each dyad, WTC values for each clusters during elaborated feedback, the example part of elaborated feedback and resting was computed by first averaging WTC matrix (time × frequency × channel) along the time dimension during the corresponding time periods, and then averaging WTC matrix along the frequency and channel dimensions based on the cluster index, and then converted into Fisher *z*-values. Finally, for each cluster, we ran step-by-step linear mixed models including a fixed effect of task (elaborated feedback vs. rest or example part of elaborated feedback vs. rest), plus random effects of learner and instructor identity.

Duration was entered in the model to control for its potential effect. Multiple comparisons were corrected using the false discovery rate (FDR) method (Benjamini & Hochberg, 1995) to calculate *corrected p* values.

**Granger causality analysis**

To estimate direction and magnitude of information flow between instructors and learners on brain regions corresponding to the identified clusters (i.e., Cluster 1, Cluster 2), granger causality analysis (GCA) was conducted via the Multivariate Granger Causality Toolbox (Barnett & Seth, 2014). GCA could quantify how well the history of one time series (e.g., X) predicts the current status of another time series (e.g., Y) with the consideration of how much Y is predicted by its own previous history. Put simply, X is said to G-cause Y if the past of X contains information that helps predict the future of Y over and above information already in the past of Y. To note, the data included only the period of elaborated feedback within each trial (noting that one full trial consisted of the question period, answer period, elaborated feedback period and confidence period). Since the focus of the current study was to investigate the instructor-learner brain activity during feedback, especially elaborated feedback, we did not perform GCA on the entire interaction period. During the period of elaborated feedback, instructors were supposed to speak while the learners were supposed to listen. Actually, learners were not forbidden to speak and they spoke some words in 13 out of 240 trials (i.e., 24 subjects*10 trials). Considering the potential influence of who was speaking on the information flow (Dai et al., 2018) between instructors and learners, those trials involving learners speaking were discarded before the GCA. Specifically, the preprocessed signals of each channel during the period of elaborated feedback was first converted into z-scores using the mean and standard deviation of the preprocessed signals during the rest, and then averaged within each cluster for each dyad. Next, for each cluster, the granger causality from instructor to learner (I→L) and from learner to instructor (L→I) were calculated for each dyad. Based on Bayesian Information Criterion (Schwarz, 1978), the model order (indicating how far back in time the model incorporates data and is measured in units of time steps) was fixed as 10 (10 Hz, 1 second) for all dyads.

Next, we used a one-sample *t*-test to respectively examine whether the values of I→L and L→I granger causality were significant. Besides, a paired-samples *t*-test was adopted to compare the values of bidirectional granger causality. Multiple comparisons were corrected using the false discovery rate (FDR) method (Benjamini & Hochberg, 1995) to calculate *corrected p* values.

#### Supplementary Results

##### Temporal patterns of instructor-learner synchronization

To investigate whether instructor-learner neural synchronization for identified clusters exhibited temporal patterns, i.e. the learners' brain activity synchronized with instructors' with some delay or the opposite, time shifts (1–14 s, step = 1 s) were added to compare the WTC during elaborated feedback or the example part of elaborated feedback vs. resting based on linear mixed models. It was revealed in Figure S4 that, with time shifts, WTC during elaborated feedback for Cluster 1 and Cluster 2 or example part of elaborated feedback for Cluster 3 and Cluster 4 remained significantly larger than that during resting ($F$s >8.04, *corrected p*s < 0.05).

##### Bidirectional information flow during providing and receiving elaborated feedback

It was shown in Figure S5 that during providing and receiving elaborated feedback, the one-sample *t*-test revealed values of granger causality from instructor to learner (I→L) and from learner to instructor (L→I) were both significantly larger than zero on CH42 and CH45 (I→L: $M \pm SD$, 0.01 ± 0.01, $t(23) = 9.75$, *corrected p* < 0.001; L→I: 0.01 ± 0.00, $t(23) = 7.91$, *corrected p* < 0.001) and CH05, CH06 and CH10 (I→L: 0.01 ± 0.00, $t(23) = 8.28$, *corrected p*

= 0.007; L→I: 0.01 ± 0.00, $t(23) = 8.13$, *corrected* $p < 0.001$). The paired-samples *t*-test revealed no significant difference between values of I→L and L→I granger causality on CH42 and CH45 ($t(23) = 1.29$, $p = 0.211$) nor CH05, CH06, and CH10 ($t(23) = 0.24$, $p = 0.812$).

**Instructor-learner neural synchronization during elaborated feedback predicts learning performance**

Next, we tested whether frontoparietal synchronized neural activity associated with elaborated feedback could predict learning performance. A SVR was trained on trial-by-trial △WTC associated with elaborated feedback respectively on Cluster 1 and Cluster 2 to predict learners' accuracy on the post-learning test relative to pre-learning test. It was revealed in Figure S6a that trial-by-trial △WTC on Cluster 1 could successfully predict out-of-sample learners' relative accuracy on the transfer measure ($r = 0.61$, $R^2 = 37.21\%$, $p = 0.002$) but not on the retention measure ($r = 0.24$, $R^2 = 5.76\%$, $p = 0.262$); trial-by-trial △WTC on Cluster 2 could not predict learning performance (*r*s < 0.10, *p*s > 0.05).

Moreover, when time shifts were added to re-perform the prediction analyses based on trial-by-trial △WTC associated with elaborated feedback on Cluster 1, the prediction accuracy on the transfer measure was significant when instructors' brain activity preceded learners' by 1–14 s and when learners' preceded the instructors' by 1–14 s (*corrected p*s < 0.05, Figure S6b). With time shifts, the prediction accuracy on the retention sub-test remained insignificant (*corrected p*s > 0.05, Figure S6b).

**Table S1**

*Experimental material*

| No. | Term | Definition | Examples |
|---|---|---|---|
| 1 | Availability heuristic | The tendency to estimate the likelihood that an event will occur by how easily instances of it come to mind. | 1. Ming and Wang worked together on a project, each putting in about half of the amount of time and effort required to finish. At the end, they have to decide who should get the most credit. Ming and Wang both claim that they did a majority of the work, probably because it is easier for them to remember their own experiences working on the project.<br>2. Find a couple and talk with them separately. If you ask the percentage of housework they have undertaken, you will receive interesting answers. The sum of their percentages will be greater than 100%. |
| 2 | Mere exposure effect | The phenomenon whereby the more people are exposed to a stimulus, the more positively they evaluate that stimulus. | 1. In a study, foreign subjects were presented with a Chinese character every two seconds, and each Chinese character was presented one, two, five, ten, or twenty-five times. Then, participants were asked to evaluate whether these Chinese characters were good or bad. As a result, it was found that the Chinese characters with more repetitions were more positively evaluated.<br>2. In one study, four equally attractive women silently attended a 200-student class for zero, 5, 10, or 15 class sessions. At the end of the course, students were shown slides of each woman and asked to rate each ones attractiveness. Students rated the women who had attended class more often as more attractive than the women who attended class less often. |
| 3 | Door-in-the-face-technique | A strategy to increase compliance based on the fact that refusal of a | 1. Ming received a telephone call from a college alumni association asking him to show his loyalty by contributing ￥1000. When he apologetically declined, the caller was sympathetic and then |

| | | | |
|---|---|---|---|
| | | large request increases the likelihood of agreement with a subsequent smaller request. | asked whether he could contribute ￥500. And if not ￥500, how about ￥200? Richard agrees to donate ￥200.<br>2. The researchers asked college students to spend two years as a voluntary counselor in a juvenile correctional institution. This is a troublesome job, and almost all college students declined. Then, the researchers put forward a small request for college students to guide the teenagers to the zoo for one time, and 50% of the people accepted the request. However, when the researchers made the latter request directly, only 16.7% of the college students agreed. |
| 4 | Fundamental attribution error | The tendency to believe that another person's behavior is due to his/her disposition and to underestimate the impact of situations on his/her behavior. | 1. When we ask the staff of the University Admissions Office for help, if the staff is indifferent, we will think that he is an unfriendly person. However, in fact we ignore the fact that the staff actually received many complaining students and thus become indifferent.<br>2. In a simple quiz game, the researchers randomly assigned the subjects as the questioner and answerer. The former asked the latter to answer some difficult questions. It was found that in this case, both the answerer and other bystanders rated the questioner as smarter, but in fact they ignored the advantage given by the role of the questioner. |
| 5 | Hindsight bias | The tendency, once an event has occurred, to overestimate one's ability to have foreseen the outcome. | 1. Some students will pat the thighs after the teacher announces the correct answer, "I know this is the choice!"<br>2. When a highly regarded basketball team loses in a huge upset, you will hear many fans claim, "I knew they were overrated and vulnerable." |
| 6 | Counterfactual thinking | A tendency to imagine alternative events | 1. It is common for the families and friends of accident victims to have endless "If only" thoughts about the accident. "If only I hadn't delayed him by |

| | | | |
|---|---|---|---|
| | | or outcomes that might have occurred but did not. | talking about …", "If only I hadn't insisted that he drive back tonight …" "If only I'd made a different plane reservation for her …" and so on.<br>2. At the end of the Olympic Games, when athletes accept medals, silver medalists are often less happy than bronze medalists. In an interview with the silver medalists, they sometimes say: "I almost won, it's too bad." |
| 7 | Deindividuation | The loss of a person's sense of individuality that results in a reduction of normal constraints against deviant behavior. | 1. Researchers have found that soldiers who hide their identities before entering the war. For example, drawing patterns on their faces and bodies, are more likely to slaughter and torture captives than soldiers who do not hide their identities.<br>2. Drivers who are less visible in convertibles with their tops up will be more likely to honk their horns at cars that fail to proceed immediately at green lights than will people who are more visible driving convertibles with their tops down. Those with tops up honk quicker, longer, and more frequently. |
| 8 | Social facilitation | A process whereby the presence of others enhances performance on simple tasks but impairs performance on more complex or unfamiliar tasks. | 1. When supervisors want their employees to complete challenging tasks, they should give employees some privacy.<br>2. If you have a good understanding of the content of the study, then reviewing with other students can help you better remember these knowledge points. However, if you do not have a good grasp of what you have learned at the beginning, and you need to learn and consolidate further, you had better choose to study alone. |
| 9 | Foot-in-the-door-technique | A strategy to increase compliance, based on the fact that agreement with a small | 1. Imagine that you work for the local animal shelter. Your goal is to increase the number of people who are willing to adopt a dog from the shelter. To maximize your success, you should first ask people if they would be willing to wear a button that says, "Adopt a dog today." A couple of weeks later, you |

| | | | |
|---|---|---|---|
| | | request increases the likelihood of agreement with a subsequent larger request. | should then ask these people to adopt a dog themselves.<br>2. Randomly interviewed some housewives and asked them to hang a small signboard on their windows. Most housewives happily agreed. After a period of time, I visited these housewives again and asked them to put a sign that was not only large but not beautiful in the courtyard. As a result, more than half of the housewives agreed. |
| 10 | Representativeness heuristic | The tendency to judge the likelihood that a target belongs to a category based on how similar the target is to typical members of the category. | 1. Mr. Zhang is a shy man. He is passionate about poetry and likes to visit art museums. People usually think of Mr. Zhang as a literary worker rather than a farmer.<br>2. If ASKED TO JUDGE the probability of flips of a coin yielding the sequence–H T H H T H–people will judge it as higher then they will if asked to judge the sequence–H H H H T H. Thus, if you expect a sequence to be random, you tend to view a sequence that “looks random” as more likely to occur. |
| 11 | Observer effect | / | / |
| 12 | Self-serving bias | / | / |

*Note.* Concept 1–10 was used for instruction and learning. Example 1 and 2 was used in elaborated feedback and transfer measure, respectively. Term 11 and 12 were additionally provided to form 12 alternatives.

**Table S2**

*The anatomical position for each channel*

| Channel | MNI coordinates x | y | z | BA | AAL |
|---|---|---|---|---|---|
| Frontal | | | | | |
| 1 | -36 | 63 | -7 | Left-BA10 | Left Middle Frontal Gyrus (Orbital) |
| 2 | -13 | 72 | -4 | Left-BA10 | Left Superior Frontal Gyrus (Orbital) |
| 3 | 15 | 71 | -3 | Right-BA10 | Right Superior Frontal Gyrus (Orbital) |
| 4 | 38 | 63 | -7 | Right-BA10 | Right Middle Frontal Gyrus (Orbital) |
| 5 | -45 | 53 | 1 | Left-BA10 | Left Middle Frontal Gyrus |
| 6 | -24 | 68 | 8 | Left-BA10 | Left Superior Frontal Gyrus |
| 7 | 2 | 68 | 9 | Right-BA10 | Right Superior Frontal Gyrus (Medial) |
| 8 | 27 | 68 | 8 | Right-BA10 | Right Superior Frontal Gyrus |
| 9 | 47 | 53 | 2 | Right-BA10 | Right Middle Frontal Gyrus |
| 10 | -35 | 58 | 19 | Left-BA10 | Left Middle Frontal Gyrus |
| 11 | -13 | 67 | 22 | Left-BA10 | Left Superior Frontal Gyrus |
| 12 | 15 | 68 | 23 | Right-BA10 | Right Superior Frontal Gyrus |
| 13 | 38 | 59 | 18 | Right-BA10 | Right Middle Frontal Gyrus |
| 14 | -45 | 42 | 27 | Left-BA10 | Left Middle Frontal Gyrus |
| 15 | -23 | 56 | 33 | Left-BA10 | Left Middle Frontal Gyrus |
| 16 | 2 | 59 | 34 | Right-BA9 | Left Superior Frontal Gyrus (Medial) |
| 17 | 26 | 57 | 33 | Right-BA9 | Right Superior Frontal Gyrus |
| 18 | 47 | 42 | 28 | Right-BA9 | Right Middle Frontal Gyrus |
| 19 | -35 | 40 | 42 | Left-BA9 | Left Middle Frontal Gyrus |
| 20 | -11 | 50 | 45 | Left-BA8 | Left Superior Frontal Gyrus (Medial) |
| 21 | 13 | 50 | 46 | Right-BA9 | Right Superior Frontal Gyrus (Medial) |
| 22 | 36 | 40 | 42 | Right-BA9 | Right Middle Frontal Gyrus |
| Left Temporoparietal | | | | | |
| 23 | -60 | -67 | -9 | Left-Fusiform (37) | Left Inferior Temporal Gyrus |
| 24 | -70 | -44 | -7 | Left-BA21 | Left Middle Temporal Gyrus |
| 25 | -70 | -17 | -8 | Left-BA21 | Left Middle Temporal Gyrus |
| 26 | -52 | -81 | 8 | Left-BA19 | Left Middle Occipital Gyrus |
| 27 | -65 | -59 | 12 | Left-BA39 | Left Middle Temporal Gyrus |
| 28 | -69 | -28 | 16 | Left-BA40 | Left Superior Temporal Gyrus |
| 29 | -65 | 1 | 16 | Left-BA6 | Left Postcentral Gyrus |
| 30 | -58 | -70 | 23 | Left-BA39 | Left Middle Temporal Gyrus |

| | | | | | |
|---|---|---|---|---|---|
| 31 | -67 | -44 | 30 | Left-BA39 | Left SupraMarginal Gyrus |
| 32 | -68 | -15 | 29 | Left-PrimSensory (1) | Left Postcentral Gyrus |
| 33 | -46 | -83 | 31 | Left-BA39 | Left Middle Occipital Gyrus |
| 34 | -59 | -59 | 39 | Left-BA39 | Left Angular Gyrus |
| 35 | -66 | -32 | 43 | Left-BA40 | Left SupraMarginal Gyrus |
| 36 | -62 | -4 | 40 | Left-PrimMotor (4) | Left Postcentral Gyrus |
| 37 | -49 | -71 | 46 | Left-BA39 | Left Angular Gyrus |
| 38 | -56 | -47 | 54 | Left-BA39 | Left Inferior Parietal Gyrus |
| 39 | -57 | -18 | 54 | Left-PrimSensory (1) | Left Postcentral gyrus |
| 40 | -28 | -81 | 51 | Left-BA7 | Left Superior Parietal Gyrus |
| 41 | -41 | -58 | 62 | Left-BA39 | Left Inferior Parietal Gyrus |
| 42 | -48 | -30 | 65 | Left-PrimSensory (1) | Left Postcentral Gyrus |
| 43 | -44 | -5 | 62 | Left-BA6 | Left Precentral Gyrus |
| 44 | -27 | -67 | 66 | Left-BA7 | Left Superior Parietal Gyrus |
| 45 | -34 | -42 | 69 | Left-PrimSensory (1) | Left Superior Parietal Gyrus |
| 46 | -32 | -16 | 70 | Left-BA6 | Left Precentral Gyrus |

**Table S3**

*Results of the cluster-based permutation test for elaborated feedback*

| Cluster | Frequency bin (Hz) | Channel | Statistic | *p* |
|---|---|---|---|---|
| 1 | 0.0168 | 42 | 11.54 | < 0.001 |
| | 0.0178, 0.0189, 0.0200, 0.0212, 0.0225, 0.0238, 0.0252 | 42,45 | | |
| 2 | 0.0168, 0.0178 | 6 | 6.62 | 0.005 |
| | 0.0189 | 10 | | |
| | 0.0200 | 10, 5 | | |
| | 0.0212, 0.0225, 0.0238 | 5 | | |
| Ns | 0.0225, 0.0238 | 20, 21 | 4.42 | 0.399 |

*Note.* Ns indicated non-significant clusters.

**Table S4**

*Results of the cluster-based permutation test for two parts of elaborated feedback*

| Cluster | Frequency bin (Hz) | Channel | Statistic | *p* |
|---|---|---|---|---|
| | | Example part | | |
| 3 | 0.0178, 0.0189, 0.0200, 0.0212, 0.0225, 0.0238, 0.0252 | 42, 45 | 13.69 | < 0.001 |
| | 0.0267 | 45 | | |
| 4 | 0.0150, 0.0159, 0.0168, 0.0178 | 6 | 10.61 | < 0.001 |
| | 0.0189 | 6, 10 | | |
| | 0.0200, 0.0212 | 5, 6, 10 | | |
| | 0.0225 | 5, 6 | | |
| Ns | 0.0225, 0.0238 | 20, 21 | 4.86 | 0.382 |
| | 0.0252 | 21 | | |
| | | Correct answer part | | |
| Ns | 0.178 | 15 | 4.87 | 0.375 |
| | 0.0189, 0.0200, 0.0212, 0.0225 | 6 | | |

*Note.* Ns indicated non-significant clusters.

**Table S5**

*Results of the cluster-based permutation test for simple feedback*

| Cluster | Frequency bin (Hz) | Channel | Statistic | *p* |
|---|---|---|---|---|
| Ns | 0.0189, 0.0200, 0.0212 | 15 | 4.97 | 0.358 |
| | 0.0238, 0.0252 | 11 | | |

*Note.* Ns indicated non-significant clusters.

**Figure S1**

*The resulting frequency bins based on wavelet transform coherence analysis.*

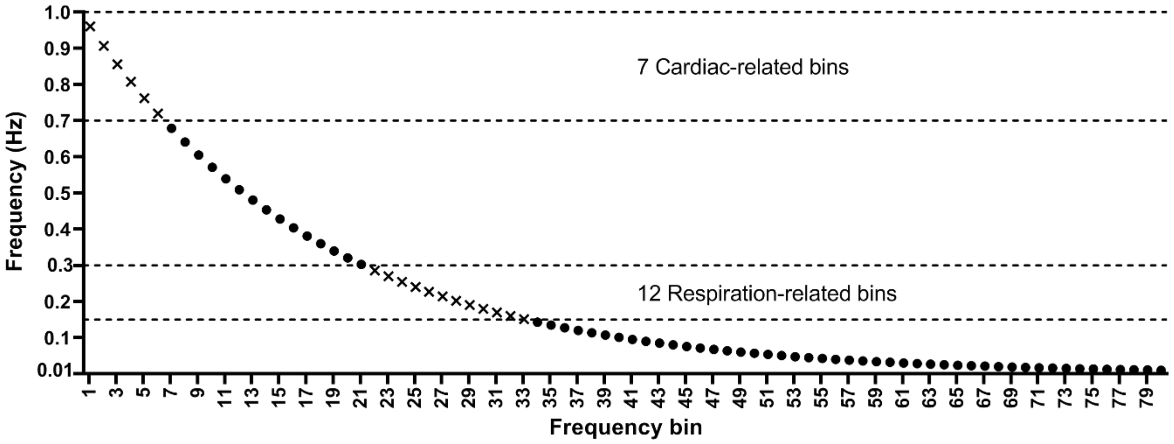


*Note.* There were totally 80 frequencies bins. 7 cardiac-related and 12 respiration-related bins (marked as crosses) were excluded.

**Figure S2**

*Schematic of permutation and data trim.*

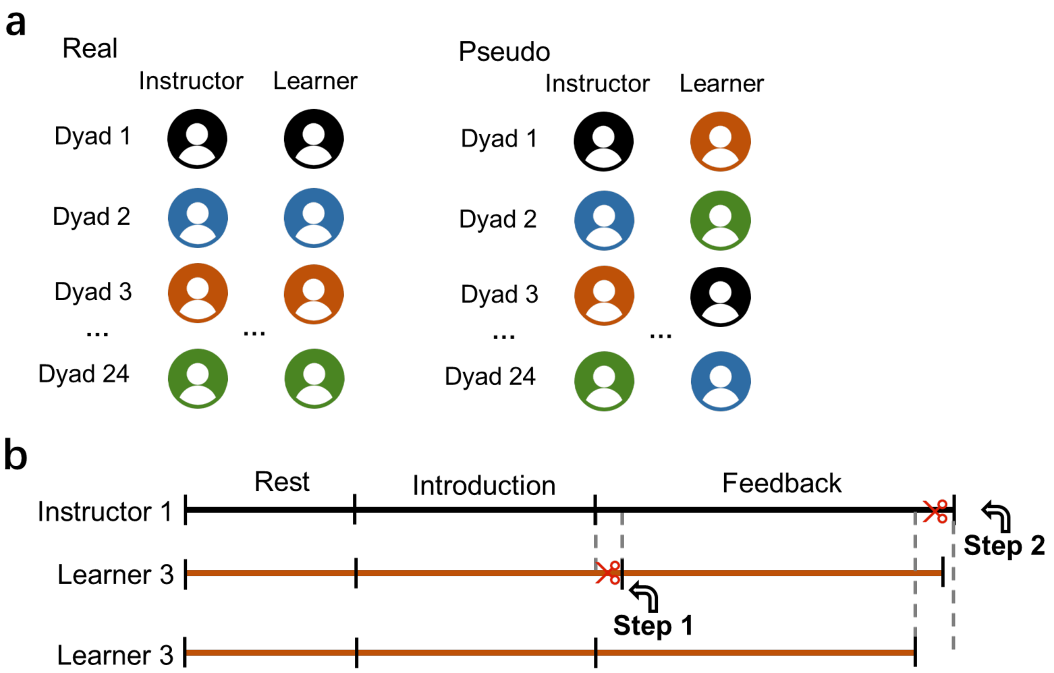


*Note.* (**a**) The permutation was conducted by randomly pairing the instructor and the learner from different original real dyads under the same condition as a pseudo dyad. (**b**) The exemplified schematic diagram of trimming longer data to match the shorter one in the pseudo dyad.

**Figure S3**

*Schematic of contrast analysis between different forms of feedback information (example vs. correct answer).*

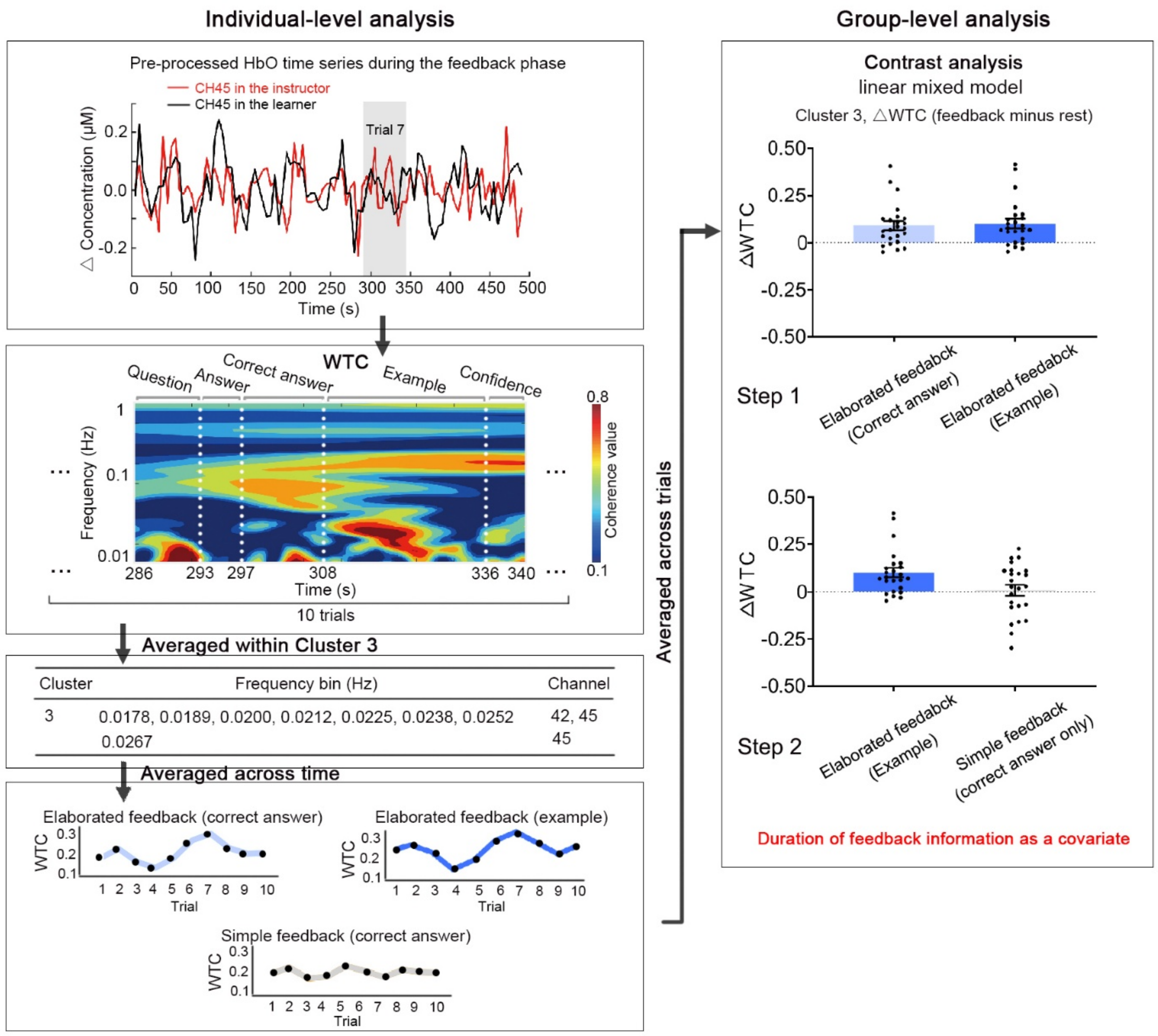


*Note.* Before entering the contrast analysis, each dyad's WTC matric was averaged within identified clusters (e.g., Cluster 3) during the 10-trial feedback. For the elaborated feedback group, the resulting time series were segmented and then averaged across time during two parts, i.e., correct answer and example, based on the recorded videos. For the simple feedback group, the resulting time series were averaged across time during the correct answer only. The contrast analysis consisted of two steps. The first was to compare trial-averaged △WTC (feedback minus rest) during the example and correct answer contained in elaborated feedback, and the second was to compare averaged △WTC during the example part of elaborated feedback and simple feedback (correct answer only) based on linear mixed models. Considering the various data length across feedback information and across dyads, duration of the feedback information was entered in the model to control for its potential effect.

**Figure S4**

*Temporal patterns of instructor-learner synchronization*

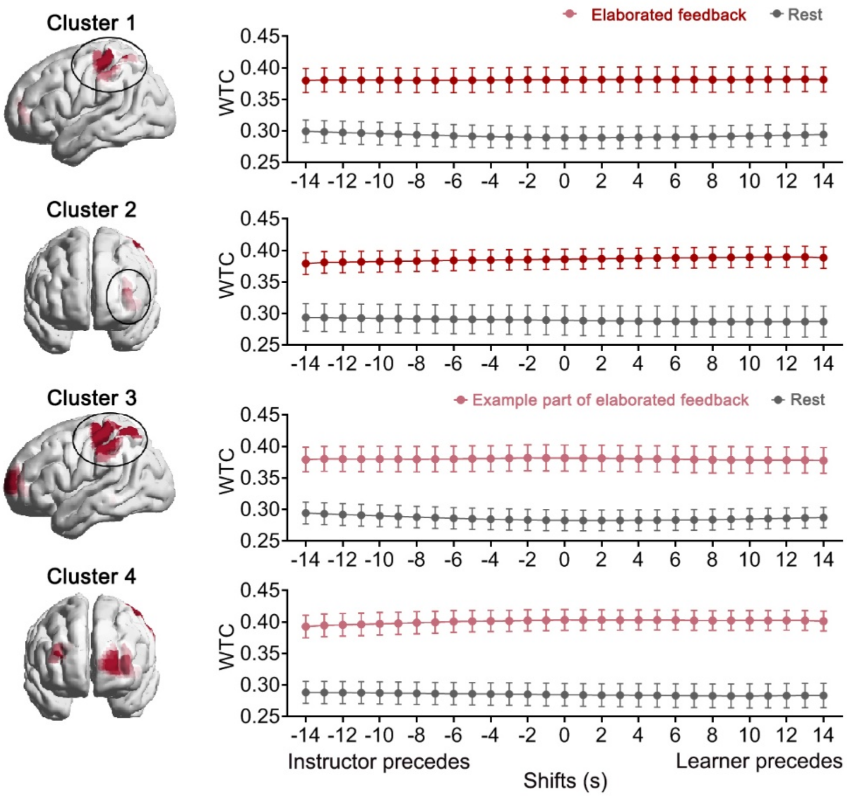


*Note.* When instructor's brain activity was first shifted forward or backward relative to the learner's by 1–14 s (step = 1 s), WTC during elaborated feedback on Cluster 1 and Cluster 2 as well as WTC during example part of elaborated feedback on Cluster 3 and Cluster 4 remained significantly larger than that during resting.

**Figure S5**

*Bidirectional information flow during providing and receiving elaborated feedback*

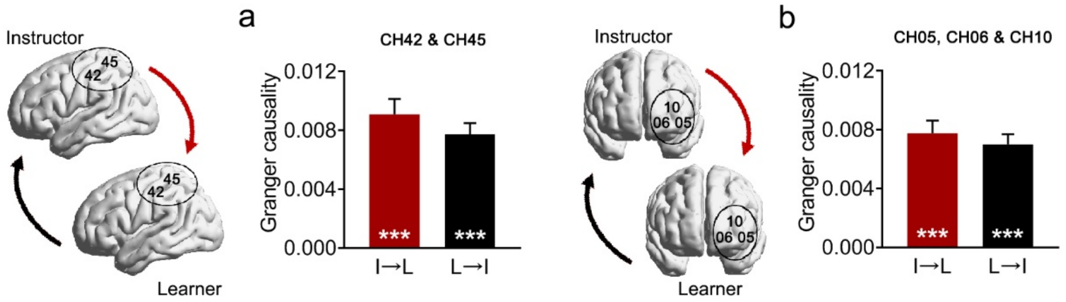


*Note.* Values of granger causality from instructor to learner and from learner to instructor were both significantly larger than zero on (**a**) CH42 and CH45 and (**b**) CH05, CH06 and CH10. I→L, from instructor to learner; L→I, from learner to instructor. ***$p < 0.001$.

**Figure S6**

*Instructor-learner neural synchronization during elaborated feedback predicts learning performance*

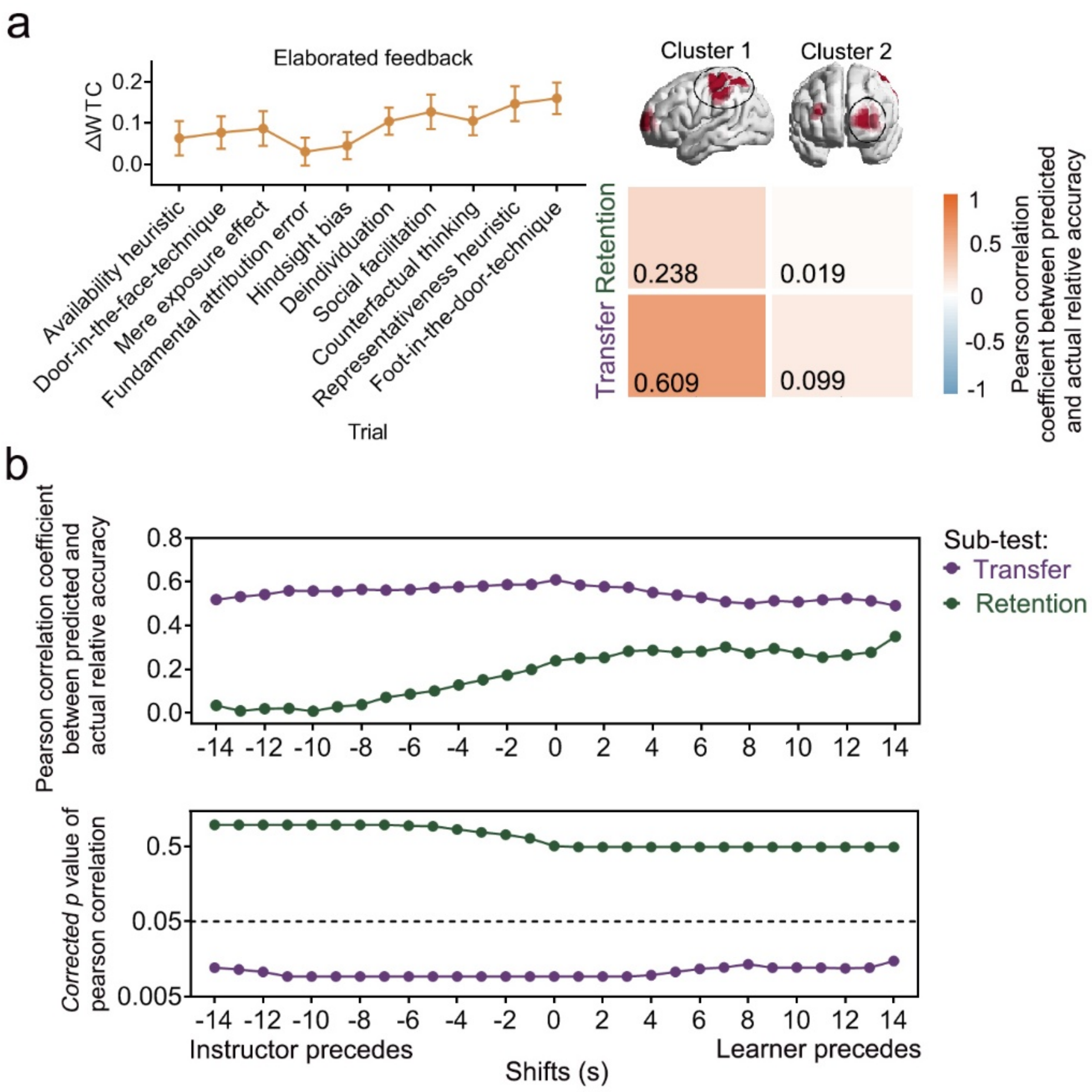


*Note.* (**a**) Trial-by-trial △WTC on Cluster 1 could successfully predict out-of-sample learners' relative accuracy on the transfer measure but not on the retention measure. Warmer colors indicate relatively higher prediction accuracy for a given cluster; cooler colors indicate relatively lower prediction accuracy for a given cluster. (**b**) The prediction accuracy for Cluster 1 on the transfer measure was significant when instructors' brain activity preceded learners' by 1–14 s and when learners' brain activity preceded instructors' by 1–14 s (-14–14, purple).
